# A new automated pipeline for Whole Genome Shotgun sequencing analysis and hazard characterization of microbial pesticides

**DOI:** 10.64898/2026.07.24.739986

**Authors:** Joao Pedro Saraiva, Valérian Lupo, Francisco Cerqueira, Stavroula Makri, Sotirios Vasileiadis, Belén Guijarro, Alexandre Bartholomäus, Costas C. Papagiannitsis, Marie Harmel, Stephan Declerck, Luc Cornet, Antonis Chatzinotas, Dimitrios G. Karpouzas, Brader Günter

**Author notes:** **Funding** This work was funded by the European Union’s Horizon Europe Research and Innovation programme in the frame of the project RATION (Risk Assessment Innovation of Low Risk Pesticides) under Grant Agreement No. 101084163. **Author contributions** JS, SV, AC, BG and DGK developed the concept and supervision of the study. JS, SV, SM, FC, VL developed the pipeline. The frontend web application was conceptually defined by JS, AB and FC and developed by Agrostis (agrostis.gr). JS, SV, VL, MH, BG and SM tested the pipeline. SV, SM, BG and KP developed the risk classification system. JS wrote the manuscript. All authors provided critical revision of the manuscript. All authors read and approved the manuscript. **Ethical statement** Not applicable.

## Abstract

Microbial pesticides are increasingly important for sustainable crop protection, yet hazard analysis and risk assessment remain challenging and require specific characterization of properties relevant to both safety and biological activity. Whole-genome sequencing (WGS) can support the identification of microorganisms at high resolution and characterize potential hazards associated with infectivity, pathogenicity, antimicrobial resistance, and toxic metabolite production. However, the routine use of WGS in this context requires accessible, reproducible, and interpretable workflows. Here, we present a publicly available web-based workflow for WGS-supported hazard analysis of microbial biocontrol agents. The workflow accepts assembled genomes as well as short-read, long-read and hybrid sequencing data, and performs genome quality assessment, taxonomic assignment, genome annotation, AMR detection, mobile-element screening, pathogenicity prediction and secondary-metabolite analysis, and compiles the results into an HTML report. For bacterial agents, we implement a transparent rule-based risk-classification module integrating taxonomic identity, PathogenFinder2 predictions, CARD/RGI resistance evidence, WHO priority taxa, and medically important antimicrobial categories. Secondary-metabolite assessment combines antiSMASH with local BLAST searches and EFSA-aligned identity/coverage thresholds to support product-level interpretation of biosynthetic gene clusters. Comparison with the currently used MOpS workflow developed by EFSA demonstrates the added benefits of the proposed workflow for a guided hazard analysis of microbial pesticides. The workflow is intended as a community-accessible pre-assessment tool that complements regulatory platforms by improving transparency, reproducibility, and early identification of potential hazards in microbial biocontrol candidates, envisioned to make WGS an integral part of the risk assessment of microbial pesticides.

**Key points:**

- The RATION-GUI provides a public, reproducible workflow for whole-genome sequencing– supported hazard characterization of bacterial and fungal microbial pesticides.
- The workflow integrates genome quality, taxonomy, annotation, antimicrobial resistance, pathogenicity-related evidence, mobile elements, and secondary-metabolite potential.
- Domain-specific outputs are translated into structured hazard indicators, evidence summaries, and recommended follow-up actions without replacing expert regulatory judgment.
- Case studies demonstrate how the workflow improves transparency, traceability, and interpretation of genomic evidence for microbial pesticide risk assessment.

## INTRODUCTION

The regulatory approval and authorization process for microbial pesticides in the European Union (EU) is governed by Regulation (EC) 1107/2009, a framework that has been built and applied to synthetic pesticides. This has long been recognized as a significant barrier to the introduction of microbial pesticides into the EU market (Frederiks & Wesseler, 2019; Robin & Marchand, 2019; Villaverde et al., 2014). While the AGRIFISH Council introduced a plan in 2016 to accelerate their adoption, the number of microbial products available in the EU remains substantially lower than in other global markets (Balog et al., 2017; *European Commission (2020a) Guidance on the Approval and Low-Risk Criteria Linked to “Antimicrobial Resistance” Applicable to Microorganisms Used for Plant Protection in Accordance with Regulation EC No. 1107/2009*, n.d.). Efforts to harmonize regulations, such as those undertaken by the OECD, and recent regulatory updates from the European Commission, aim to address these challenges (*European Commission (2020a) Guidance on the Approval and Low-Risk Criteria Linked to “Antimicrobial Resistance” Applicable to Microorganisms Used for Plant Protection in Accordance with Regulation EC No. 1107/2009*, n.d.; *European Commission (2020b) Guidance on the Risk Assessment of Metabolites Produced by Microorganisms Used as Plant Protection Active Substances in Accordance with Article 77 of Regulation EC No. 1107/2009*, n.d.).

Currently, the primary microbial pesticides available in the EU market are 73. Amongst them, only 30 have been classified as low-risk products (LRPs), including bacterial strains (e.g. *Bacillus velezensis, B. subtilis, Pasteuria nishizawae*), fungal strains (e.g. *Clonostachys rosea, Trichoderma atroviride, Isaria fumosorosea*), and viral strains (e.g. mild pepino mosaic virus). Emerging solutions, such as microbial consortia, phages and protists,s are anticipated to enter the market soon, with four phage products pending authorization in the EU. While microbial consortia have seen extensive applications in medicine, bioengineering and as biostimulants, their potential in crop protection has only more recently garnered attention (Zhang et al., 2018).

Relevant guidance documents recently released by the European Commission have addressed the most important issues in the risk assessment of microbial pesticides, including their infectivity, pathogenicity, production of toxic secondary metabolites (*European Commission (2020b) Guidance on the Risk Assessment of Metabolites Produced by Microorganisms Used as Plant Protection Active Substances in Accordance with Article 77 of Regulation EC No. 1107/2009*, n.d.), and the potential transferability of antibiotic resistance (AMR) traits (*European Commission (2020a) Guidance on the Approval and Low-Risk Criteria Linked to “Antimicrobial Resistance” Applicable to Microorganisms Used for Plant Protection in Accordance with Regulation EC No. 1107/2009*, n.d.). Additionally, the European Food Safety Agency (EFSA) has published guidelines on performing Whole Genome Sequence (WGS) analysis for microorganisms when required for risk assessment (RA) (Authority (EFSA), 2021).

Although new regulatory documents address certain hazard-specific issues, comprehensive guidance for microbial pesticide RA remains limited. Tools like WGS analysis and advanced bioinformatics could address challenges directly relevant for microbial strains, such as AMR dispersal and toxic metabolite production, streamlining the potential planning of risk assessment and data generation of new microbial pesticides.

EFSA has developed the Microorganisms Pipelines Service (MoPS) (https://www.efsa.europa.eu/sites/default/files/corporate_publications/files/amp2224.pdf, page 88), a platform destined only for internal use by regulatory bodies and designed to harmonize WGS-based re-analysis of microorganisms in regulatory assessments. MoPS provides standardized pipelines for bacteria, yeasts, filamentous fungi and viruses, including quality control, taxonomic identification and detection of traits of concern. However, MoPS was developed as a general-purpose regulatory platform for microorganisms across multiple sectors, including food, feed and genetically modified microorganisms, and primarily functions as an evidence aggregation and reporting system. Although highly valuable for standardized characterization, its outputs still require substantial expert interpretation on a case-by-case basis, particularly for microbial pesticides where risk assessment increasingly focuses on properties such as infectivity, pathogenicity, antimicrobial resistance relevance and the production of toxic secondary metabolites. Recent European guidance and legislation have further clarified expectations regarding these endpoints and their role in microbial pesticide risk assessment, increasing the need for workflows capable not only of reporting genomic evidence, but also of supporting its transparent and reproducible interpretation in a risk assessment context.

To address this need, we developed a publicly available, user-friendly WGS-based hazard analysis workflow and graphical user interface for microbial biocontrol candidates. By integrating risk-relevant endpoints into a reproducible and guided pre-assessment framework, the workflow aims to support the early identification of potential hazards and facilitate the evaluation of microbial biocontrol agents. The workflow complements existing regulatory infrastructures by enabling transparent pre-screening of bacterial and fungal candidates, automated extraction of WGS-derived hazard indicators, and generation of an HTML report summarizing the main findings, evidence tables, and potential follow-up tasks.

## MATERIALS AND METHODS

The workflow for hazard analysis of bacterial or fungal biocontrol strains using WGS data is executed using a Web-server Graphical User Interface (GUI) which was implemented using JavaScript, PHP, and MySQL on a standard LAMP stack and is publicly available at https://agrostis.gr/ration-ra-gui. Current resources available to the web server consist of 22 CPUs with 120 GB of RAM and one GPU with 80 GB of RAM, with the possibility of future expansion.

### Workflow for hazard analysis of bacterial potential low-risk pesticides using WGS

The hazard analysis of bacterial pesticides using WGS data consists of five or six stages depending on whether raw sequence data is provided. If the latter is true, then additional steps of sequence data pre-processing and assembly are executed. Subsequent steps all follow the same stages: quality control, taxonomic assignment, genome annotation, virulence and pathogenicity prediction, assessment of secondary metabolism and risk classification (Figure 1). The whole procedure, including the analyses performed and data extraction, is designed to comply with the EFSA statement on requirements for WGS analysis for microbes proposed as biocontrol agents (Authority (EFSA), 2021) presented in an HTML report. Unless stated otherwise, default values are used for all tools integrated in the pipeline. Singularity containers were built for each of the tools incorporated into the workflow. For each run, the workflow records tool versions, command-line parameters, database paths or versions where available, container image paths, and image identifiers. These metadata are automatically included in the HTML report, providing run-level provenance rather than relying solely on a generic methods description. All tools, including versions, are listed in Table 1.

**Table 1.** Tools incorporated in the pipeline, including versions and a short description of their functions.

| Tool | Version | Description |
| --- | --- | --- |
| Filtlong | v0.2.1 | Quality filtering and trimming of long-read sequencing data |
| Unicycler | v0.5.0 | Hybrid assembly tool combining short- and long-read data for high-quality bacterial genome assemblies |
| CheckM | v1.2.3 | Assesses genome completeness and contamination |
| GTDBtk | v2.3.2 | Classifies genomes using the Genome Taxonomy Database for accurate taxonomic assignment |
| ISEscan | 1.7.2.3 | Detects insertion sequences (IS elements) within genomes |
| StarAMR | v0.10.0 | Identifies antimicrobial resistance genes using a curated database |
| RGI | v6.0.3 | Detects and annotates resistance genes (typically linked with the CARD database) |
| DFAST | v1.3.1 | Provides rapid annotation for prokaryotic genomes |
| fastp | v0.23.2 | Fast and efficient quality control and preprocessing of sequencing reads |
| VirSorter | v2.2.4 | Identifies viral sequences within microbial genomes or metagenomes |
| BLAST | v2.16.0 | Performs sequence similarity searches against nucleotide or protein databases |
| antiSMASH | v7.1.0 | Detects and annotates secondary metabolite biosynthetic gene clusters |
| Circleator | v1.0.2 | Visualizes circular genomes and genomic features |
| PathogenFinder2 | v3.11.14 | Prediction of human pathogen |

**Figure 1.**
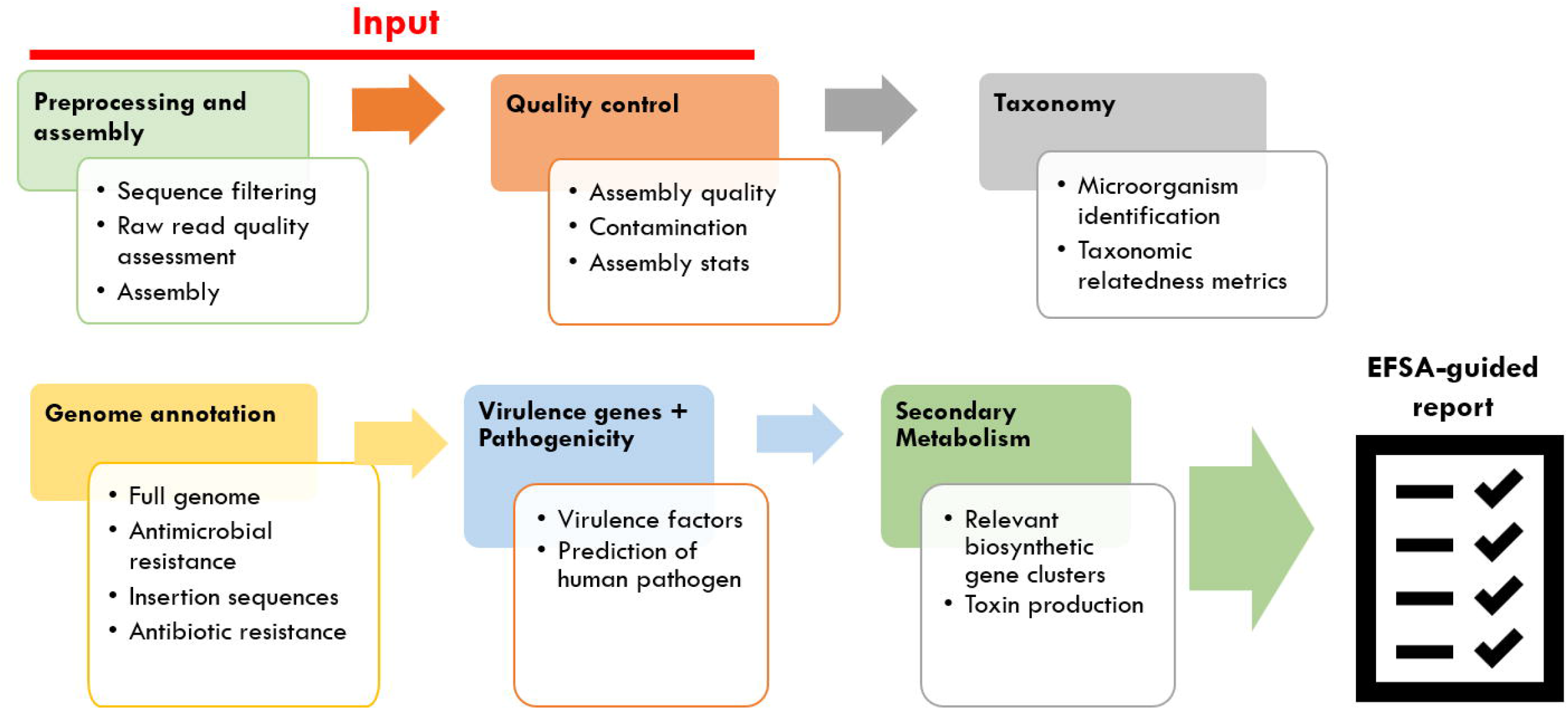
WGS-based hazard analysis workflow for bacteria. If short paired-end reads are submitted, their preprocessing (e.g., adapter removal and read trimming) and quality are assessed using FastP v0.23.2. If long reads are provided, filtering and quality assessment is performed using Filtlong v0.2.1. Assembly is performed using Unicycler v0.5.0. Quality of assembled reads is assessed using CheckM v1.2.3. Taxonomy is assigned using GTDBtk v2.3.2. Genome annotation is carried out by DFast v1.3.1. AMR prediction is done by RGI v6.0.3 and StarAMR v.0.10.0. Potential transferability of AMR is assessed by identifying insertion sequences in the genome, which, in turn, is performed using ISEscan v.1.7.2.3. Phages are identified using VirSorter2 v2.2.4. Secondary metabolism is assessed by coupling the AntiSMASH v7.1.0 results with local BLAST v2.16.0 local alignments against the target genome. Lastly, a risk classification is assigned according to the rules outlined in the section *Bacterial Hazard classification system*.

### Bacterial Hazard classification system

The bacterial risk-classification module was designed as a transparent prioritization layer rather than as a final regulatory conclusion. Its purpose is to translate WGS-derived evidence into an interpretable screening category that helps users identify genomes requiring further expert review, phenotypic testing or contextual assessment (Figure 2). The classification therefore complements, but does not replace, regulatory expert judgement. It is currently implemented for bacteria, where curated taxonomic, pathogenicity and antimicrobial-resistance evidence can be combined in a rule-based framework.

**Figure 2.**
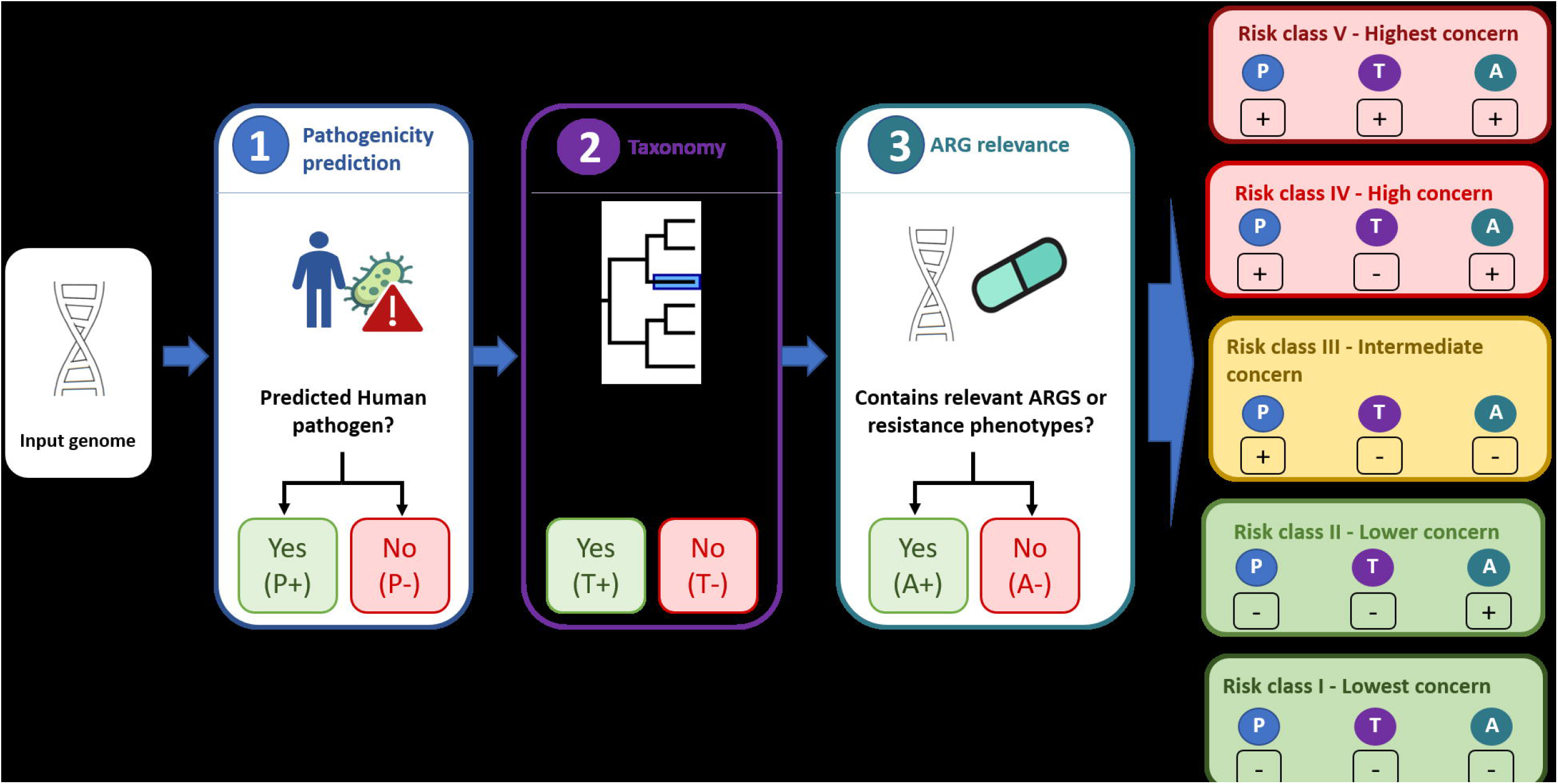
Simplified bacterial risk-classification scheme.

The proposed ranking system follows the logic of One Health risk-management priorities, WHO priority pathogen information, and European guidance on the relevance of antimicrobial resistance in microbial active substances. Microbial genomes are assigned to classes according to predicted pathogenicity to humans, taxonomic affiliation, and the presence and medically relevant predicted antimicrobial-resistance phenotypes. Pathogenicity is inferred using PathogenFinder2 (Florensa et al., 2026), taxonomic assignment is derived from GTDB-Tk (Parks et al., 2022), and resistance evidence is obtained using the CARD Resistance Gene Identifier (RGI) (Alcock et al., 2023). Antibiotic-importance categories are mapped to the accompanying WHO/WOAH/FAO and European antimicrobial-relevance tables (Supplementary files S1-S3).

For each genome, ARGs are aggregated across all detected hits and assigned to the most conservative category detected. When multiple categories are present, the highest-risk class prevails. Five bacterial classes are used, with class V representing the strongest screening flag and class I representing the lowest screening priority in the current implementation. Classes II and IV are subdivided according to the medical relevance of the affected antimicrobial category.

Risk class V is assigned to genomes when (i) PathogenFinder2 predicts human pathogenicity, (ii) the strain is classified in a WHO priority pathogen species according to GTDB-Tk (Supplementary file S1), and (iii) at least one ARG maps to a WHO priority resistance combination or to an antimicrobial category considered highly relevant for human or animal health. Within this class, two probabilistically possible ranks can be assigned: (V.a) with positive ARG identification on the verified WHO human pathogenic strain; (V.b) without ARG annotations for the WHO pathogenic strain.

Risk class IV is assigned to genomes predicted as human pathogens by PathogenFinder2 that are not classified as WHO priority taxa but carry predicted resistance phenotypes. Subclasses distinguish resistance to human-only antimicrobials (IV.a), highest-priority critically important or critically important antimicrobials used in humans and animals (IV.b), highly important or important antimicrobials used in humans and animals (IV.c), and animal-only antimicrobials (IV.d).

Risk class III is assigned to genomes that are not classified as WHO priority taxa, are predicted as human pathogens by PathogenFinder2, and do not show predicted resistance phenotypes associated with medically important antimicrobial categories in the CARD-RGI analysis.

Risk class II is assigned to genomes predicted as non-pathogenic to humans but carrying predicted resistance phenotypes. Subclasses distinguish resistance to human-only antimicrobials (II.a), highest-priority critically important or critically important antimicrobials used in humans and animals (II.b), highly important or important antimicrobials used in humans and animals (II.c), and animal-only antimicrobials (II.d).

Risk class I is assigned to genomes predicted as non-pathogenic by PathogenFinder2 and lacking predicted resistance phenotypes associated with medically important antimicrobial categories in the CARD-RGI analysis. This class should be interpreted as a low-priority WGS screening result, not as proof of absence of hazard under all product-use scenarios.

The complete list of risk categories and interpretation for bacteria is shown in Table 2.

**Table 2.** WGS-based risk categories and interpretation layers used in the bacterial workflows. Bacterial genomes are assigned to classes I–V using taxonomic information, predicted human pathogenicity, and antimicrobial resistance evidence, with subclasses for the medical relevance of the antimicrobial category. *Note: These categories are intended as transparent screening and prioritisation outputs and do not replace expert regulatory judgement*.

| Domain | Risk category | Main decision criteria | Subclassification | Interpretation |
| --- | --- | --- | --- | --- |
| Bacteria | Risk class V | Genome is assigned to a WHO priority pathogen taxon, predicted as human pathogenic by PathogenFinder2, and carries | V.a: ARGs identified on the WHO classified and PathogenFinder2 | Highest screening flag. Requires expert review, contextual assessment, |
|  |  | at least one ARG linked to a WHO priority resistance combination or to an antimicrobial category of high relevance for human or animal health. | characterised human pathogenic strain; V.b: no ARG identification on the WHO classified and PathogenFinder2 characterised human pathogenic strain | and, where relevant, phenotypic confirmation. |
| **Bacteria** | Risk class IV | Genome is predicted as human pathogenic by PathogenFinder2, is not assigned to a WHO priority pathogen taxon, and carries predicted resistance phenotypes. | IV.a: human-only antimicrobials; IV.b: highest priority critically important or critically important antimicrobials used in humans and animals; IV.c: highly important or important antimicrobials used in humans and animals; IV.d: animal-only antimicrobials. | High screening priority due to predicted pathogenicity combined with AMR evidence. |
| **Bacteria** | Risk class III | Genome is predicted as human pathogenic by PathogenFinder2 but does not carry predicted resistance phenotypes associated with medically relevant antimicrobial categories in CARD RGI. | Not subdivided | Pathogenicity-driven screening flag without medically relevant AMR evidence. |
| **Bacteria** | Risk class II | Genome is predicted as non-pathogenic to humans by PathogenFinder2 but carries predicted resistance phenotypes. | II.a: human-only antimicrobials; II.b: highest priority critically important or critically important antimicrobials used in humans and animals; II.c: highly important or important antimicrobials used in humans and animals; II.d: animal-only antimicrobials. | AMR-driven screening flag. Requires interpretation of ARG identity, genomic context, intrinsic versus acquired origin, and potential mobility. |
| **Bacteria** | Risk class I | Genome is predicted as non-pathogenic by PathogenFinder2 and lacks predicted resistance phenotypes associated with medically important antimicrobial categories in CARD RGI. | Not subdivided | Lowest screening priority in the current bacterial implementation. This does not prove absence of hazard under all use scenarios. |
| **Bacteria** | Evidence interpretation | Results are interpreted together with taxonomic resolution, genome quality, tool outputs, antimicrobial or metabolite relevance, and biological context. | Confirmed, probable, or putative evidence may be distinguished where appropriate. | The workflow provides a transparent pre-assessment and prioritization framework, but final conclusions require expert judgement and may require phenotypic, chemical, or regulatory follow-up. |

### Detection of secondary metabolism

Microbes can produce bioactive compounds which may be toxic to humans and/or non-target organisms. Thus, it is critical to determine which metabolites may be produced by the target biocontrol organism. To this end, the genome of the microorganism is analysed with antiSMASH, resulting in one or more predicted biosynthetic gene clusters (BGCs). For each predicted BGC with a reported similarity to a known cluster greater than 0, the corresponding MIBiG record is queried, and the genes/products from the known reference cluster are retrieved. These sequences are then used to generate BGC-specific BLAST databases, which are searched against the query genome by local sequence alignment.

The resulting BLAST alignments are filtered to highlight hits with sequence identity ≥80% and coverage of the MIBiG/antiSMASH reference subject sequence ≥70%, mirroring the threshold logic used in EFSA/MoPS-style reporting of genes of concern. When multiple local BLAST alignments occur for the same query–subject pair, subject coverage is calculated after merging the aligned subject intervals, thereby reducing the impact of repeated short local High-scoring Segment Pairs (HSPs). For each BGC-specific BLAST table, the report also indicates how many predicted reference products passed these thresholds relative to the total number of products in the corresponding known cluster, allowing users to distinguish between near-complete and partial cluster similarity. Products were labelled as “CRUCIAL” when the corresponding MIBiG/antiSMASH gene annotation contained the term “biosynthetic”. This rule was used as a conservative proxy for core biosynthetic genes within the reference BGC, while transport, regulatory, and accessory genes were retained in the full BLAST output but were not labelled as crucial.

### Workflow for risk assessment of fungal biocontrols using WGS

The risk assessment (RA) of fungi using WGS data consists of four or five stages depending on whether raw sequence data is provided. When raw data are available, an additional stage involving data pre-processing and assembly is performed (i.e., short-read, long-read, or hybrid assembly). Afterwards, the same general stages are applied: genome quality control, taxonomic support, genome annotation, and assessment of secondary-metabolite biosynthetic potential (Figure 3). Unlike the bacterial workflow, a formal fungal risk-classification module is not yet implemented, because fungal pathogenicity and mycotoxin risk require different curated evidence layers and interpretation rules. The whole procedure, analyses performed, and data extraction comply with the EFSA statement on requirements for WGS analysis for fungi proposed as biocontrol agents (7) and are presented in an HTML report. Unless stated otherwise, default values are used for all tools integrated in the pipeline. Singularity containers were built for each of the tools incorporated into the workflow. All tools, including versions, are listed in Table 3.

**Table 3.** Tools incorporated in the fungal pipeline including versions and a short description.

| Tool | Version | Description |
| --- | --- | --- |
| MaSuRCA | v4.1.0 | Genome assembler pipeline combining short- and/or long-read data into high-quality contiguous sequences |
| Flye | v2.9-b1778 | Genome assembler for long reads generated by PacBio and Oxford Nanopore Technologies |
| EukCC | v2.1.1 | Assesses eukaryotic genome completeness and contamination |
| BUSCO | v5.4.5 | Evaluates genome completeness based on conserved single-copy orthologs and can assist in taxonomic inference |
| Barrnap | v0.9 | Identifies ribosomal RNA genes in assembled genomes |
| Forty-Two | v0.213470 | Phylogenomic tool that integrates sequences into existing alignment while controlling for orthology and contamination |
| BRAKER | v3.0.8 | Annotates eukaryotic genomes and predicts gene structures |
| antiSMASH | v7.1.0 | Detects and annotates secondary metabolite biosynthetic gene clusters |
| Palantir | v0.251630 | Toolbox for post-processing and parsing antiSMASH report |
| BLAST | v2.9.0+ | Performs sequence similarity searches against nucleotide or protein databases |
| Bio-MUST-Core | v0.252040 | Perl module providing utilities for sequence data processing |
| Bio-MUST-Drivers | V0.252830 | Perl module providing wrappers to execute external bioinformatic tools |

**Figure 3.**
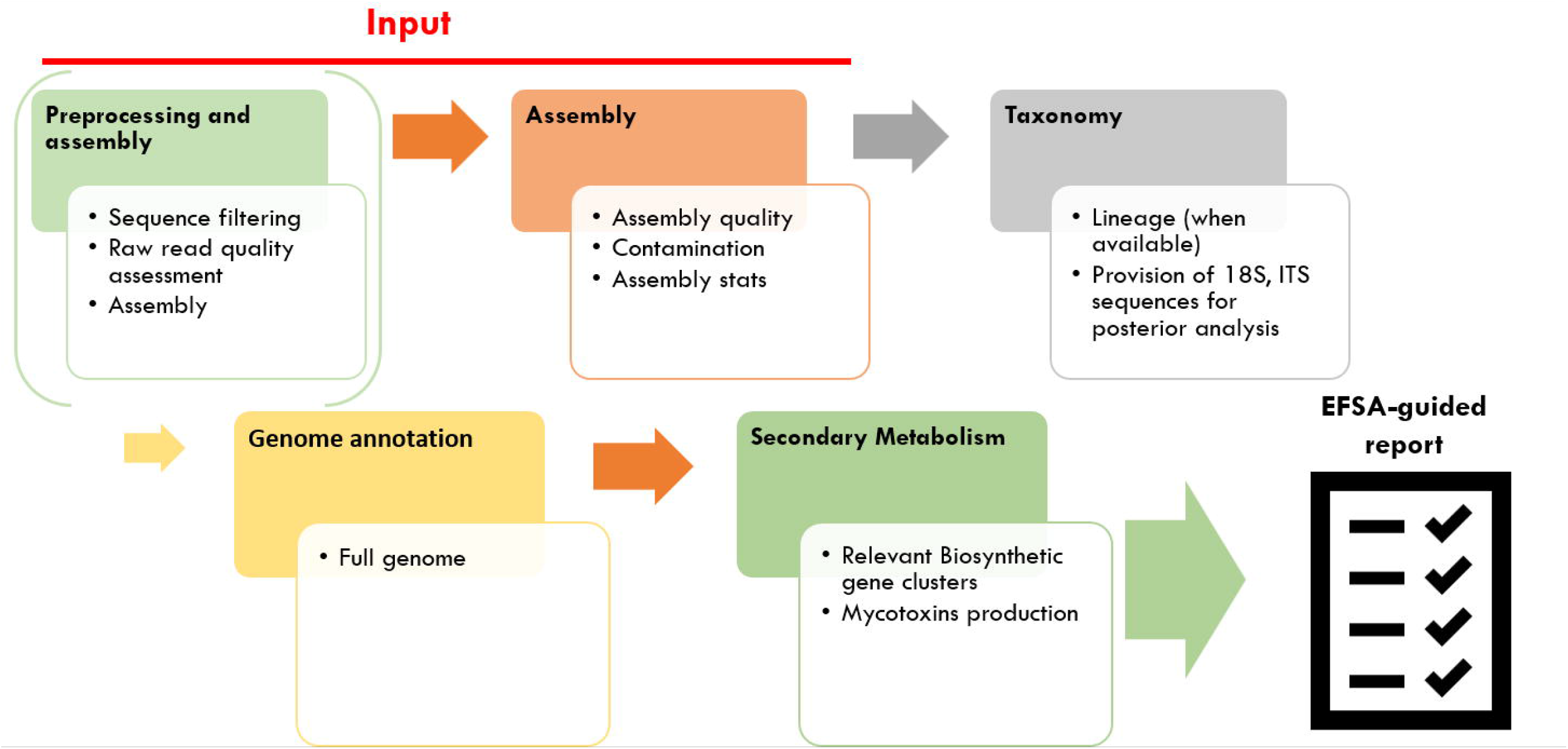
WGS-based fungal risk assessment workflow. Raw read pre-processing and genome assembly are performed using MaSuRCA v4.1.0 for Illumina paired-end reads, and hybrid assemblies combining Illumina short reads with either PacBio or Nanopore long reads. When only PacBio or Nanopore long reads are provided, filtering and quality assessment are performed using Flye v2.9-b1778. Quality control of assembled genomes is performed using EukCC v2.1.1 and BUSCO v5.4.5. Taxonomy can be inferred from the lineage assigned by BUSCO, when available, and/or further determined by BLAST analysis of predicted ribosomal RNA (rRNA) gene cluster sequences against the NCBI rRNA database. The 18S, 5.8S, and 28S rRNA sequences are identified using Barrnap v0.9, while full ITS (ITS1—5.8S—ITS2) sequences are identified using Forty-Two v0.213470 through similarity searches against the NCBI fungal ITS database (latest version downloaded at each run). Genome annotation is carried out by BRAKER v3.0.8 with Fungi OrthoDB v12 (Tegenfeldt et al., 2025) as protein evidence. Secondary metabolism is assessed by antiSMASH v7.1.0 and parsed with the Palantir v0.251630 Perl modules.

### Mycotoxin Database Generation

During this study, two databases were used to identify relevant secondary metabolites in fungi. The first corresponds to a list of MIBiG accession numbers (Supplementary files S4 and S5) from repository v4.0 (downloaded on 19 March 2025), for which the main product name matches either a mycotoxin compound listed in the MycoCentral database (downloaded on 26 June 2024) or an antibiotic compound listed in NCBI Pathogens database (downloaded on 22 May 2025) or in Antibiotic Combination DataBase (ACDB) (Lv et al., 2022).

The second database consists of BGC proteins involved in the biosynthetic pathways of mycotoxin compounds considered as of concern by EFSA for human and veterinary health to complement the list of mycotoxins not covered by MIBiG (Supplementary files S6 and S7). The sequences were downloaded from NCBI on 12 November 2025.

### Detection of secondary metabolism

The query genome is analyzed using antiSMASH with the ‘KnownClusterBlast’ option enabled, allowing BLAST-based similarity comparisons between all predicted BGCs and experimentally characterized gene clusters from the MIBiG database. antiSMASH then reports matches to known clusters with high-ranking scores (see https://docs.antismash.secondarymetabolites.org/modules/clusterblast/#ranking-system) and provides gene-level metrics, including percentage of identity, coverage, bit-score, and e-value. Finally, the Palantir toolbox, together with the first database (list of MIBiG accessions), is used to filter matches corresponding to mycotoxins and antibiotics.

In parallel, all predicted proteins from the query genome are compared against the second database (proteins involved in mycotoxin biosynthesis) using a best-hit BLAST identity search with an e-value threshold set to 1e-10.

### Fungal Hazard classification system

For fungi, a formal risk classification is not yet implemented; instead, the workflow supports structured assessment of genome quality, taxonomy, annotation, and secondary metabolite biosynthetic potential.

### Web-server Graphical User Interface (GUI)

The user is only required to provide the sequence file(s), a name for the project, the type of data being submitted (e.g., assemblies, long reads, short reads), and the domain of the candidate (prokaryotic or fungi). When creating a new project, the user first selects the domain of the candidate genome (Figure 4a). A new window then opens, where the user provides the remaining information required to configure and launch the analysis (Figure 4b).

**Figure 4.**
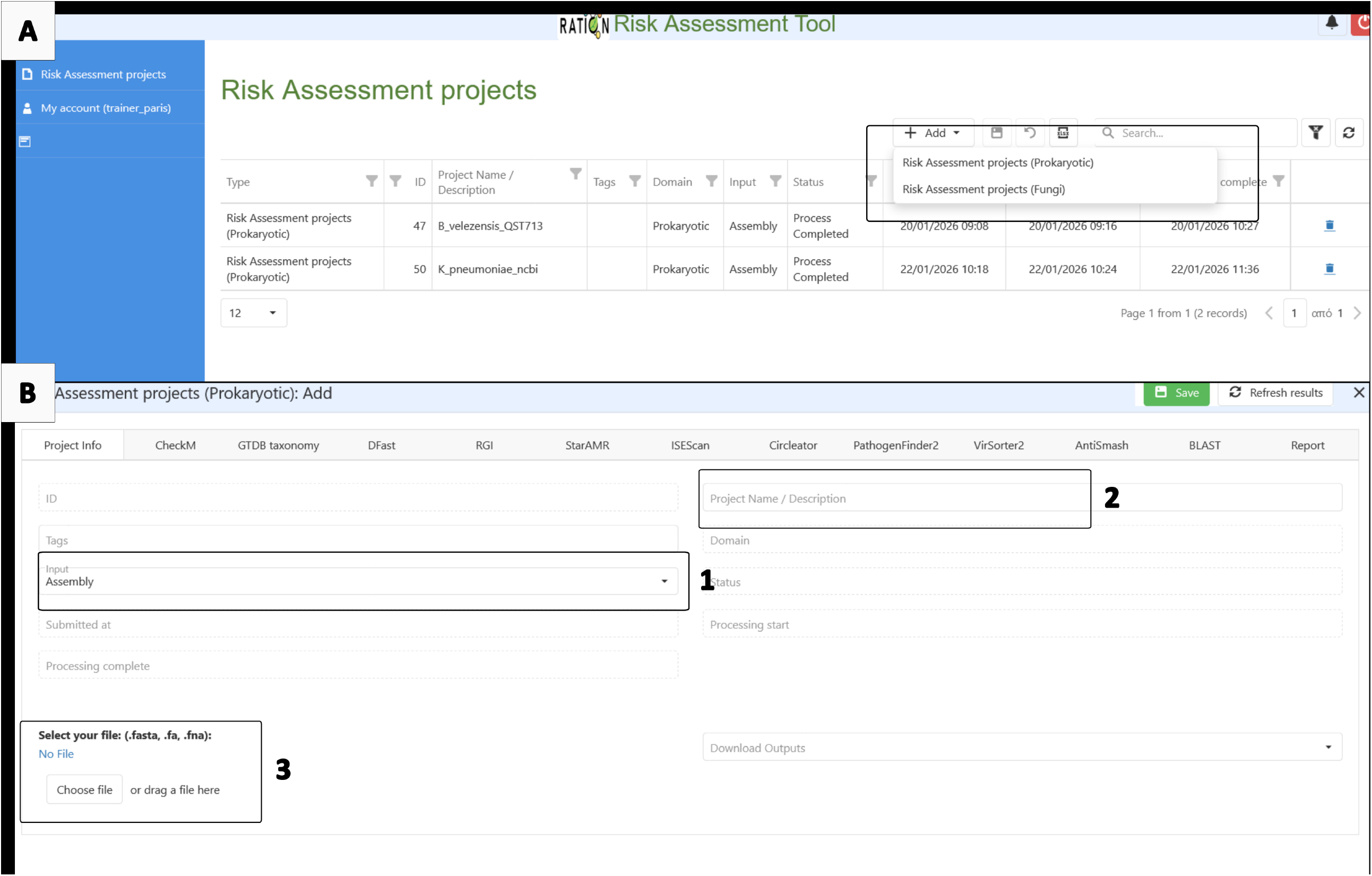
(A) Selection of candidate genome domain; (B) User input for a bacterial candidate: 1-Type of sequence data; 2 - Project name; 3 - File upload.

An overview of all projects is available in the main window, where the user can track the progress of the workflow and access the results (Supplementary file S8). Once a project is processed, the user can directly access and download the individual results via the different displayed tabs (Supplementary file S9a). The user also has the option to download a compressed folder with all results and store them locally (Supplementary file S9b). Users can also access the HTML report directly, which compiles the information according to EFSA guidelines and summarizes all relevant information collected from all tools (Figure 5).

**Figure 5.**
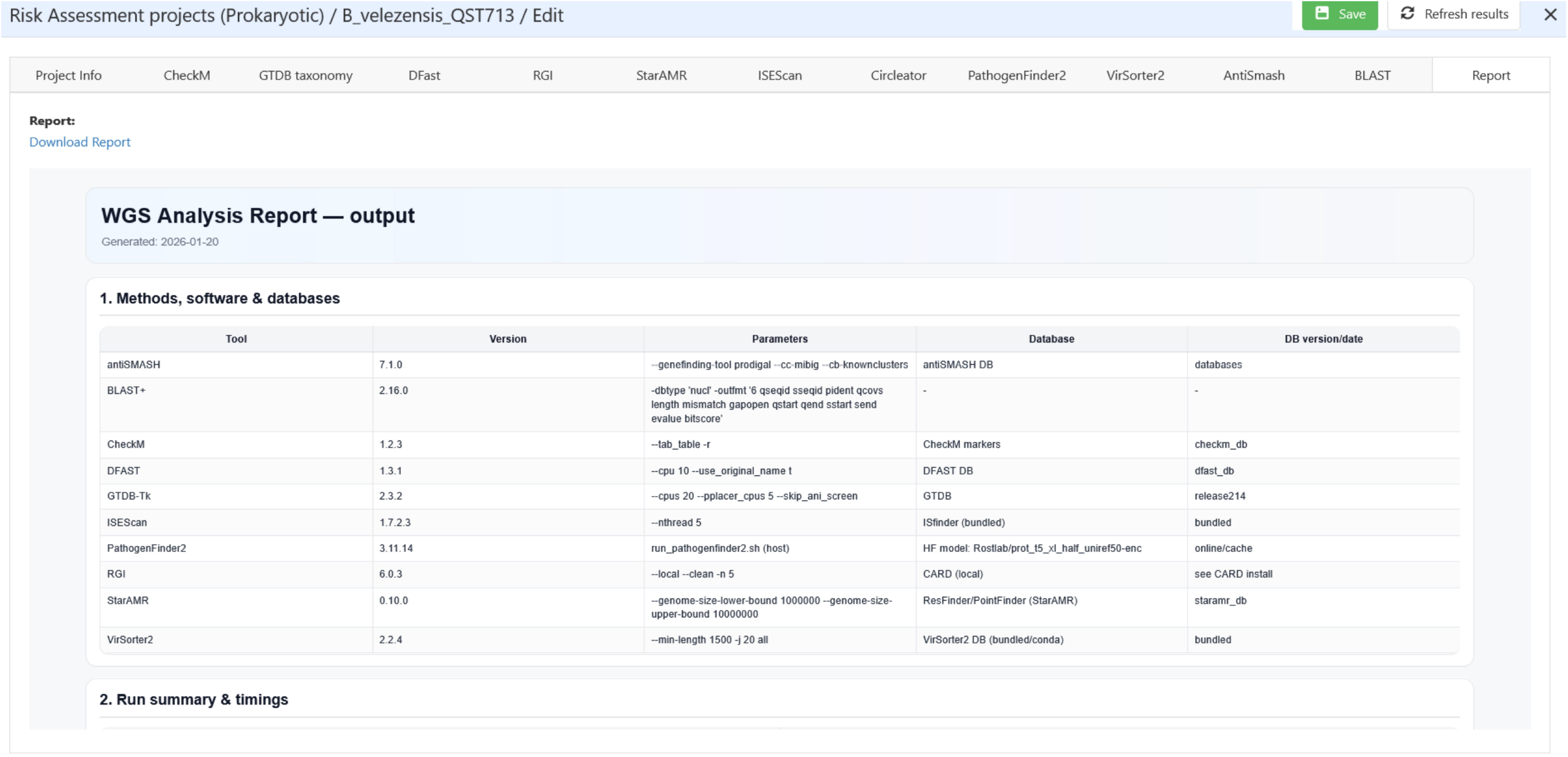
HTML report.

## Results and Discussion

### Genomes used as case studies

To showcase the use of the workflow, we processed and analyzed through our workflow the genomes of a bacterium (*Bacillus velezensis* QST713 – Accession number GCF_003073255.1) and a fungus (*Beauveria bassiana* HN6 - Accession number GCA_014607475.1) used as plant protection products using the web server.

### Case study – *Bacillus velezensis* QST713

The applicability of the proposed workflow was evaluated using *Bacillus velezensis* QST713, a commercially relevant microbial biocontrol strain with a documented history of agricultural use and broad-spectrum antagonistic activity against plant pathogens. QST713 constitutes a particularly relevant case study because *Bacillus* species are commonly characterised by multiple modes of action, including rhizosphere colonisation, antimicrobial activity and induction of plant defense responses, while also representing a microbial group frequently associated with regulatory uncertainty regarding secondary metabolites, pathogenicity, persistence and antimicrobial resistance (AMR) (Dėlkus et al., 2026; Poulaki & Tjamos, 2023).

All results mentioned in the case study are accessible via the compressed output folder (Supplementary file 5). The processing of the complete workflow for the already assembled genome required ∼71 minutes, with VirSorter2, GTDB-tk, and ISEScan being the major bottlenecks in processing time. Assembly quality (Completeness of 99.59% and 0% contamination) was above the required threshold (Completeness ≥ 94% and contamination ≤ 5%). The taxonomy was also correctly assigned with an Average Nucleotide Identity (ANI) of 97.99% (EFSA requires more than 94% for species relatedness). Additional steps can be taken by the user, such as confirming if the species is mentioned in a risk class by other technical documents (e.g., Technical Rules for Biological Agents (TRBA) of the Federal Institute for Occupational Safety and Health (BAuA)).

According to the EFSA statement (2024) on the requirements for WGS analysis of microorganisms intentionally used in the food chain, a fully assembled bacterial genome should be provided. This also resolves obstacles with fragmented gene clusters for secondary metabolites with a high likelihood to occur due to the modular structure of Non-ribosomal peptide synthetases (NRPS) and polyketide synthases (PKS) (Klassen & Currie, 2012). However, in the specific case here, only a single sequence was obtained. A total of eight ARGs were obtained using RGI, but only one of them (*clbA* from the Cfr 23S ribosomal RNA methyltransferase family) showed an identity value greater than 80%, as suggested as a threshold in EFSA 2024. Only four mobile genetic elements were found related to two Insertion sequence families (IS*21* and IS*3*). A total of five double-stranded DNA phages were obtained with mixed viral and cellular scores, which suggest integrated prophages (Wishart et al., 2023). Follow-up tasks for the user may include assessing the genomic context of the AMR hits by linking their position to the mobile genetic elements, which in this case does not show mobile elements in the vicinity of *clbA*; the nearest detected IS element was more than 550 kb away. In addition, a follow-up includes assessment of the intrinsic nature of the ARG. The common occurrence (detected in >460 completed genomes of *B. velezensis* with tblastn at NCBI on 03.04.2026) of the ARG *clbA* confirms the intrinsic nature of this ARG in *B. velezensis*.

PathogenFinder2 classified the genome as human non-pathogenic. In the bacterial risk-classification module, the genome was assigned to risk class II.a, reflecting a non-pathogenic prediction combined with the presence of at least one AMR-like determinant associated with medically relevant antimicrobials. This classification should be interpreted as a screening flag for targeted follow-up rather than as evidence of unacceptable risk. The risk class was driven by the RGI hit to *clbA*, a Cfr-family 23S rRNA methyltransferase detected with high sequence identity. The VirSorter2 and ISEScan outputs should therefore be retained as supporting evidence documenting the lack of co-localization between the AMR-associated locus and mobile genetic elements.

antiSMASH predicted 14 biosynthetic gene cluster (BGC) regions, of which 10 showed similarity to known MIBiG reference clusters and were therefore evaluated by local BLAST-based comparison against the candidate genome. After applying the EFSA/MoPS-aligned filtering criteria of sequence identity ≥80% and merged subject-sequence coverage ≥70%, 8 of these 10 known-cluster regions yielded reportable region-specific BLAST tables. Regions 1.3 and 1.11 were evaluated but did not yield full hits passing both final thresholds; lower-coverage or multi-fragment similarities, where present, were retained separately in the partial-review output rather than interpreted as confirmed full product-level matches.

The strongest product-level conservation was observed for several biocontrol-associated BGCs. In particular, the difficidin, macrolactin H, bacillaene, fengycin, and bacilysin-associated regions showed complete or near-complete product-level support under the final thresholds. Difficidin, macrolactin H, bacillaene, fengycin, and bacilysin each showed all reference products passing the applied identity and subject-coverage criteria, while surfactin and subtilin showed partial but substantial product-level conservation. In contrast, the locillomycin-like region showed more limited similarity, with only a subset of reference products passing the final thresholds.

The secondary-metabolite findings should be interpreted in terms of both potential efficacy and potential hazard. The presence of intact biocontrol-associated clusters, such as lipopeptide-related clusters, may support the expected biological activity of the strain, but does not by itself establish the amount of metabolite produced under product-relevant conditions. Conversely, detection of similarity to known clusters should not automatically be interpreted as toxicological concern unless cluster completeness, core biosynthetic genes, compound identity, production conditions and exposure are considered. Furthermore, this case also illustrates an important limitation of automated BGC interpretation: neighbouring or overlapping clusters may be represented as a single antiSMASH region, and compound-level prediction generally remains at the metabolite-family level, requiring manual inspection of gene content, module architecture, substrate predictions and literature.

Automated outputs should be interpreted together with targeted expert review, including taxonomic confirmation, assessment of AMR relevance and genomic context, secondary-metabolite interpretation, and, where appropriate, phenotypic or chemical confirmation. Detailed recommendations for these follow-up procedures are provided to users in a dedicated guidance document (Supplementary file S10).

Overall, the *B. velezensis* QST713 case study shows how the bacterial workflow converts a complete genome into an interpretable pre-assessment report. The genome fulfilled quality and taxonomic requirements, was classified as human non-pathogenic, and contained only one high-identity AMR hit above the applied threshold. The combined interpretation of AMR results, ISEScan outputs, and VirSorter2 predictions suggested that this hit should be prioritized for contextual assessment rather than treated as immediate evidence of transferable resistance. The secondary-metabolite module reduced a large number of raw BGC alignments to a smaller set of high-confidence matches, while also exposing interpretation challenges such as partial cluster similarity and neighbouring BGCs. While this case study demonstrates the utility of automated WGS screening, expert review of AMR context, intrinsic resistance evidence, BGC completeness, literature-supported safety and, where relevant, chemical or phenotypic confirmation is still required.

The complete output folder of the *B. velezensis* QST713 case study is available as a compressed folder in the supplementary materials (Supplementary file S11).

### Comparison with MoPS output

To place the RATION output in the context of the currently used EFSA Microorganisms Pipelines Service (MoPS), we compared the exported output folders generated for the *B. velezensis* QST713. Both MoPS and RATION captured the high-identity AMR hit; however, RATION presented the evidence in an organized, contextualized, and translated into a reproducible hazard-analysis report.

MoPS provided a rich evidence set, including AMRFinderPlus, ResFinder, ABRicate/hAMRonization, plasmid and prophage-related outputs, and reference-comparison files. However, the output was distributed across many tool-specific folders and required substantial manual interpretation to connect AMR evidence, mobile-element context, prophage calls, taxonomy and secondary-metabolite information. In contrast, RATION preserved access to individual tool outputs but added a consolidated HTML report, run-level provenance, EFSA-aligned filtering logic, explicit follow-up tasks and a rule-based bacterial risk-classification layer. This makes the output more directly usable for early-stage microbial pesticide pre-assessment, where transparency, auditability and consistent interpretation are as important as detection.

Considering the AMR and virulence factor results presented in Table 4, clear differences were observed between the RATION workflow and the EFSA MOP pipeline, reflecting their distinct objectives in microbial risk assessment. The RATION workflow identified a broader set of putative antimicrobial resistance determinants, including tet(45) (74.62% identity), FosBx1 (64.23%), BcI (63.4%), and qacJ (46.84%). These genes were associated with resistance mechanisms such as antibiotic efflux, antibiotic inactivation, and target alteration. However, most of these hits exhibited moderate to low sequence identity, suggesting that they represent homologous resistance-associated genes rather than confirmed acquired resistance determinants.

**Table 4.**
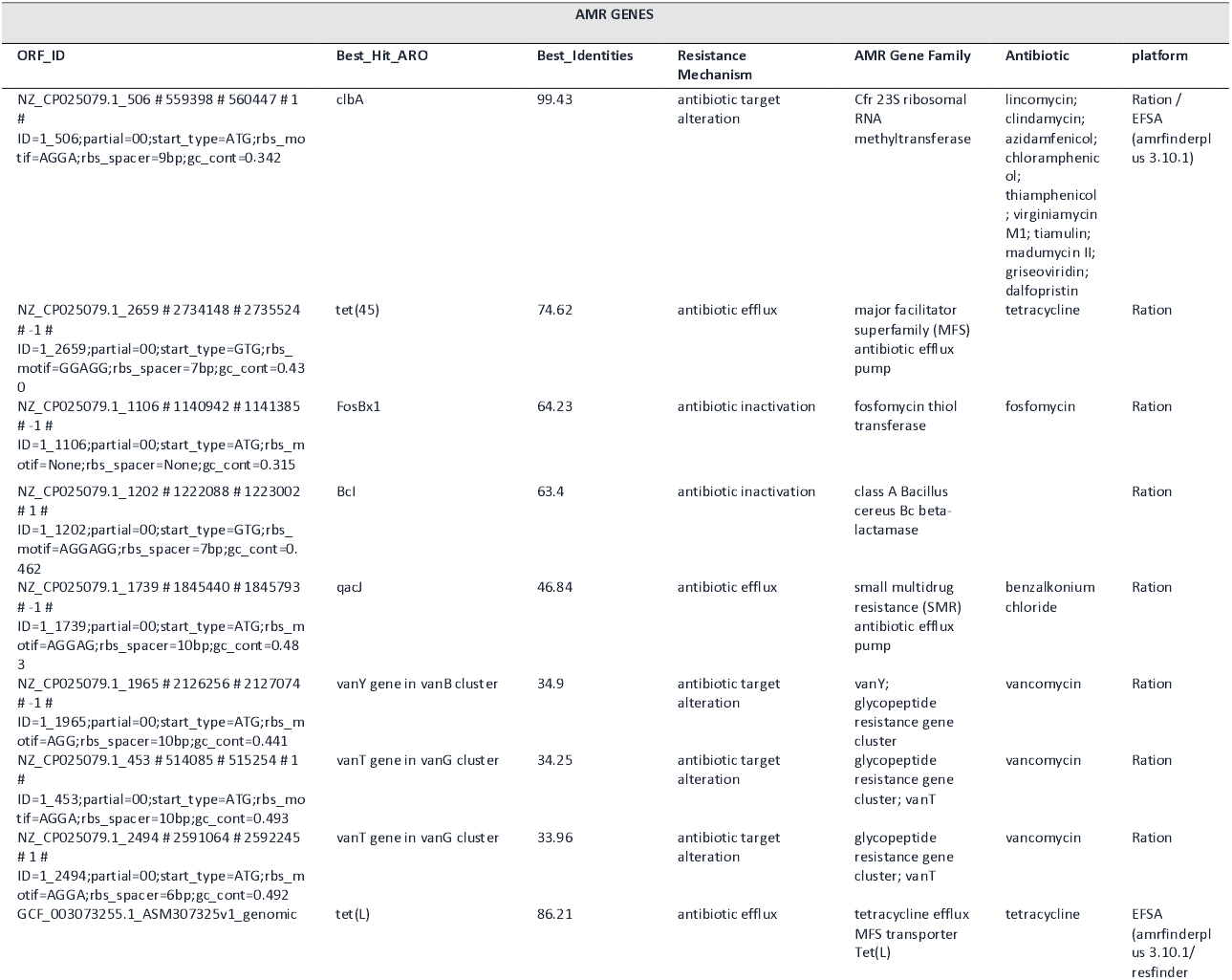
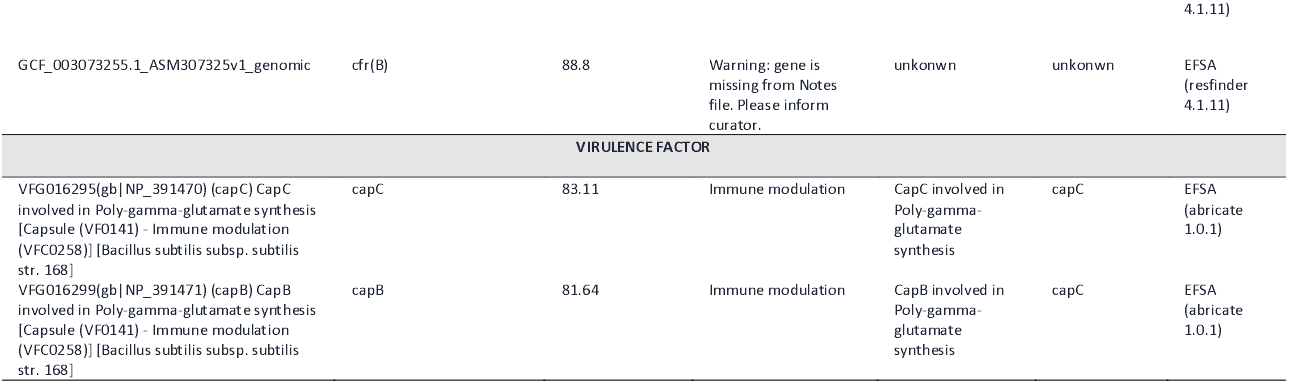
Comparison between MoPS and RATION in the identification of AMR genes and virulence factors for *B. velezensis* QST713.

In contrast, the EFSA MOP pipeline produced a more conservative AMR profile, retaining only tet(L) (86.21% identity) and cfr(B)/clbA (99.43% identity by AMRFinderPlus; 88.8% identity in ResFinder) as higher-confidence findings. The detection of tet(L) is consistent with a tetracycline efflux mechanism commonly reported in environmental Gram-positive bacteria and represents the strongest evidence for a potential AMR determinant in the strain. Likewise, the identification of a cfr-like ribosomal RNA methyltransferase is noteworthy because this gene family can confer reduced susceptibility to several classes of antimicrobials, including phenicols, lincosamides, pleuromutilins, streptogramins, and oxazolidinones. Importantly, this determinant was detected by both RATION and EFSA-based analyses, making it the most robust AMR-related finding in the dataset. Nevertheless, the warning associated with the ResFinder annotation indicates that additional manual verification would be required before drawing definitive conclusions regarding its functionality and regulatory significance.

The EFSA workflow additionally identified the virulence-associated genes capB and capC, both involved in poly-γ-glutamate capsule biosynthesis and classified within immune-modulation functions. Although these genes are traditionally included in virulence factor databases, their occurrence in *Bacillus velezensis* should not be interpreted as evidence of pathogenicity. Instead, they are more likely associated with environmental fitness, biofilm formation, stress tolerance, and rhizosphere colonization, traits that contribute to the ecological success of beneficial plant-associated Bacillus species. This interpretation is consistent with the absence of known toxin genes, the non-pathogenic classification obtained through pathogenicity prediction, and the long history of safe use of QST713 as a microbial biocontrol agent.

From a risk assessment perspective, the combined results suggest that *Bacillus velezensis* QST713 possesses a limited number of high-confidence AMR-associated determinants, primarily represented by tet(L) and the putative cfr-like methyltransferase, while the majority of RATION-exclusive hits should be considered exploratory evidence requiring further validation. The hybrid interpretation provided by combining both workflows offers a more balanced assessment than either approach alone. The EFSA pipeline provides regulatory confidence through stringent filtering and curated databases, whereas the RATION workflow increases sensitivity and supports early identification of potentially relevant resistance-associated genes that may warrant additional investigation. Consequently, the principal safety considerations for QST713 are not related to pathogenicity but rather to the potential functionality and transferability of the confirmed AMR determinants and their ecological relevance under conditions of use. Such a hybrid framework reduces the risk of overlooking biologically meaningful signals while avoiding overinterpretation of weak homology-based matches, thereby providing a more comprehensive basis for microbial pesticide risk assessment.

Table 5 summarizes the main differences observed between the two exported output structures and highlights the added value of RATION as a guided, public and regulator-oriented pre-assessment workflow rather than a replacement for expert review.

**Table 5.** Comparison of MoPS and RATION output structures.

| Dimension | MoPS | RATION | Added value of RATION |
| --- | --- | --- | --- |
| **Primary purpose** | Broad microorganism characterization and evidence aggregation across regulatory contexts. | Microbial pesticide-focused hazard analysis and pre-assessment. | Focuses interpretation on endpoints relevant to microbial active substances. |
| **Output organization** | Extensive but fragmented tool-specific folders, requiring manual navigation and synthesis. | Tool-specific folders plus a consolidated WGS HTML report and summary tables. | Improves readability, extraction of key findings and hand-over to assessors. |
| **AMR evidence** | AMRFinderPlus, ResFinder, ABRicate and hAMRonization provide broad AMR evidence layers. | RGI and StarAMR provide resistance evidence. | Improvement lies mainly in contextual integration. |
| **Transferability context** | Mobile-element and plasmid-related outputs are available but require manual linkage to AMR loci. | ISEScan, AMR coordinates and report-level follow-up tasks support inspection of AMR genomic context. | Makes the distinction between intrinsic/background resistance and potentially transferable resistance easier to assess. |
| **Secondary metabolism** | Provides evidence layers that still require manual interpretation at product or cluster level. | Combines antiSMASH with local BLAST, identity/coverage thresholds and region-specific tables. | Improves product-level interpretation of BGC similarity and flags partial versus near-complete matches. |
| **Provenance and auditability** | Contains raw outputs but less directly exposes run-level methods and parameters in a single report. | Records tool versions, parameters, database paths/versions, logs and run timings in reportable form. | Strengthens reproducibility and regulatory traceability. |
| **Risk-oriented synthesis** | Primarily reports evidence requiring expert synthesis. | Adds a transparent bacterial risk-classification module. | Converts raw WGS evidence into a guided screening outcome while preserving expert-review requirements. |

### Case study – *Beauveria bassiana HN6*

All results mentioned in the case study are accessible via the compressed output folder (Supplementary file S6). The processing of the complete workflow for the assembled fungal genome required ∼71 minutes, with BRAKER being the major bottleneck in processing time. Assembly quality using EukCC reported a completeness value of 100% and 0 contamination. Genome completeness was also assessed using BUSCO v5.4.5 with the hypocreales_odb10 dataset (n = 4,494), yielding 96.7% complete BUSCOs (96.6% single-copy, 0.1% duplicated), 1.0% fragmented, and 2.3% missing (EFSA requires BUSCO values greater or equal to 95%). The assembly comprised 12 gapless contigs totalling 37.12 Mb, consistent with a near chromosome-level fungal genome assembly which also satisfies EFSA requirements of fewer than 1000 sequences/contigs for fungal genomes. These results support the use of the assembly for secondary-metabolite screening, as high completeness and low contamination are required to minimize false absence or false presence of genes of concern. As a follow-up taxonomic confirmation step, the rRNA and ITS sequences extracted by the workflow should be queried against appropriate NCBI reference databases using BLASTn/megablast, and the identity of the 18S, ITS, 5.8S and 28S regions should be compared to determine the most plausible taxonomic assignment.

AntiSMASH predicted 54 Biosynthetic Gene Clusters (BGCs) across 10 of the 12 contigs, confirming the extensive secondary-metabolite potential expected for a filamentous fungal biocontrol candidate. The antiSMASH overview included several matches to known clusters, encompassing clusters related to Beauveria-associated metabolites such as beauvericin, bassianolide, tenellin/desmethylbassianin, oosporein and beauveriolide-type compounds. The best-hit BLAST search against the curated mycotoxin-related protein database produced 832 entries, of which 201 were located within predicted BGCs. Sixteen BLAST hits showed ≥80% identity; however, only nine of these high-identity hits were also located within predicted BGCs, mainly corresponding to beauvericin/bassiatin and one beauveriolide-associated hit. These results support the presence of known *B. bassiana* secondary-metabolite pathways while also showing that raw protein-level identity searches require careful interpretation, especially when hits occur outside predicted BGCs or correspond to isolated genes rather than complete biosynthetic pathways.

Fungal genomes tend to have a higher number of clusters with unknown identity. This is due to the fact that certain metabolites are only produced under very specific developmental conditions or due to common silencing of clusters combined with a lack of knowledge for these clusters (Keller, 2019). For the analysis of the output and follow-up tasks, in addition to all aspects discussed above for bacteria, the following fungal-specific points should be considered: (i) In most cases, the gene prediction and intron splicing are not based on transcriptome or proteome evidence. Hence, the overall identity of specific genes might be underestimated in case of a discrepancy in prediction. Care has to be taken if complete clusters are present but individual genes of the cluster have lower identity values under the threshold, and the alignment needs to be inspected manually; and (ii) Fungi are known for their potential to produce highly toxic mycotoxins. In the pipeline, a mycotoxin analysis independent of MIBiG is implemented, allowing a prediction of all critical mycotoxins (Battilani et al., 2020) in the species of interest except moniliformin, for which biosynthesis clusters are currently unknown. In the case of *Beauveria bassiana* HN6, the following critical mycotoxin clusters were not detected, and hence production of aflatoxins and their precursors sterigmatocystins, Alternaria toxins (altenuene and AAL-toxins, alternariols, altertoxins, stemphyltoxins, tentoxins and tenuazonic acids), citrinin, cyclochlorotin, enniatin, ergot alkaloids, fumonisins, ochratoxin, phomopsins, trichothecenes (including Fusarium toxins) and zearlenones can be ruled out. The Beauveria-associated metabolite clusters responsible for beauvericin, bassiatin, and beauveriolide were, however, detected.

The *B. bassiana* HN6 case study demonstrates that the fungal workflow can generate a compact but information-rich overview of genome quality, completeness, taxonomic support and secondary-metabolite potential. The assembly met quality expectations, with 12 gapless contigs, 100% EukCC completeness, no detected contamination, and 96.7% complete BUSCOs using the hypocreales_odb10 dataset. The secondary-metabolite analysis identified 54 antiSMASH regions and highlighted several known Beauveria-associated metabolites, while the curated BLAST layer provided an additional screen for mycotoxin-related biosynthetic proteins. This case study also illustrates why fungal hazard analysis cannot yet be reduced to a simple rule-based class: metabolite prediction depends on cluster completeness, gene-model accuracy, expression conditions, compound-specific toxicological relevance and product-level exposure. The fungal workflow is therefore best interpreted as a structured prioritization tool that identifies candidate clusters and proteins requiring expert review, literature evaluation and, where necessary, analytical chemistry confirmation.

The complete output folder of the *B. bassiana* HN6 case study is available as a compressed folder in the supplementary materials (Supplementary file S12).

### Concluding remarks

The proposed publicly available web-based workflow shares several analytical principles with existing WGS-based frameworks developed for microorganism characterization in regulatory settings, but extends these principles into a more transparent, interpretable, and community-accessible framework for microbial pesticide hazard analysis. The QST713 comparison with MoPS outputs illustrates that the key improvement is not simply a larger number of raw detections, but a clearer route from WGS-derived evidence to auditable interpretation.

The workflow preserves the analytical foundations expected from WGS-supported characterisation, including quality assessment, taxonomy, genome annotation, AMR screening and secondary-metabolite analysis, while adding a risk-detection-oriented interpretation layer for bacteria. By integrating taxonomic identity, PathogenFinder2 predictions, AMR evidence, antimicrobial-relevance categories and taxonomic risk information, RATION provides a guided decision-support framework that improves consistency while retaining the need for expert judgement.

The workflow also expands the assessment of pathogenicity by integrating predictive modelling approaches in addition to conventional screening of virulence-associated determinants. This feature is particularly relevant for microbial pesticides, where infectivity and unintended pathogenicity are important considerations in the characterisation of microbial active substances. Likewise, although secondary metabolite prediction remains based on established biosynthetic gene cluster detection tools, the addition of sequence identity searches and predefined identity and coverage thresholds improves the regulatory interpretability of predicted metabolites and reduces the risk of overestimating biologically irrelevant clusters.

Another important distinction is accessibility. While many existing systems remain institutionally restricted or require substantial bioinformatics expertise, the proposed workflow is implemented as a publicly accessible web-based platform, facilitating reproducibility and enabling broader use by researchers, applicants and risk assessors during early-stage hazard identification and pre-assessment activities.

Finally, unlike broader microorganism characterisation frameworks developed across multiple regulatory sectors, the proposed workflow was designed specifically for microbial biocontrol agents and microbial pesticides. This domain-specific focus enables prioritisation of endpoints that are particularly relevant for pesticide risk assessment, including pathogenicity, AMR relevance and toxic secondary metabolite production.

In conclusion, we developed a publicly accessible WGS-based workflow and GUI to support early-stage risk assessment of microbial biocontrol candidates. The workflow builds upon existing regulatory infrastructures such as MoPS by providing a transparent and user-oriented implementation for pre-submission screening. Its main added value lies in the combination of automated multi-tool analysis, containerized execution, run-level provenance, EFSA-aligned threshold reporting, secondary-metabolite interpretation, and rule-based bacterial risk classification. The generated HTML report is intended to help applicants and assessors identify potential hazards and define follow-up analyses, including phenotypic AMR testing, manual inspection of BGC completeness, assessment of ARG genomic context, and chemical confirmation of metabolite production. Future development should focus on expanding fungal risk classification, implementing additional contamination and type-strain comparison modules, and benchmarking the workflow against additional curated regulatory case studies. The workflow is designed to be upgradable and expandable in response to emerging genomic challenges in the microbial pesticide market. Future extensions could accommodate WGS analysis of phages, which are currently pending authorisation in the EU market, as well as protists, for which innovation and production platforms remain less developed.

## Supplementary data

Supplementary data includes two compressed files (.zip) with all project folders for each case study.

## REFERENCES

Alcock, B. P., Huynh, W., Chalil, R., Smith, K. W., Raphenya, A. R., Wlodarski, M. A., Edalatmand, A., Petkau, A., Syed, S. A., Tsang, K. K., Baker, S. J. C., Dave, M., McCarthy, M. C., Mukiri, K. M., Nasir, J. A., Golbon, B., Imtiaz, H., Jiang, X., Kaur, K., … McArthur, A. G. (2023). CARD 2023: Expanded curation, support for machine learning, and resistome prediction at the Comprehensive Antibiotic Resistance Database. Nucleic Acids Research, 51(D1), D690–D699. 10.1093/nar/gkac920

Authority (EFSA), E. F. S. (2021). EFSA statement on the requirements for whole-genome sequence analysis of microorganisms intentionally used in the food chain. EFSA Journal, 19(7), e06506. 10.2903/j.efsa.2021.6506

Balog, A., Hartel, T., Loxdale, H. D., & Wilson, K. (2017). Differences in the progress of the biopesticide revolution between the EU and other major crop-growing regions. Pest Management Science, 73(11), 2203–2208. 10.1002/ps.4596

Battilani, P., Palumbo, R., Giorni, P., Dall’Asta, C., Dellafiora, L., Gkrillas, A., Toscano, P., Crisci, A., Brera, C., De Santis, B., Rosanna Cammarano, R., Della Seta, M., Campbell, K., Elliot, C., Venancio, A., Lima, N., Gonçalves, A., Terciolo, C., & Oswald, I. P. (2020). Mycotoxin mixtures in food and feed: Holistic, innovative, flexible risk assessment modelling approach: EFSA Supporting Publications, 17(1), 1757E. 10.2903/sp.efsa.2020.EN-1757

Dėlkus, M., Ivanauskas, A., Žižytė-Eidetienė, M., Lukša-Žebelovič, J., Vepštaitė-Monstavičė, I., Brokevičiūtė, S., & Šimkutė, N. (2026). Bacillus Species in Agriculture: Functional Traits, Biocontrol Performance, and Regulatory Safety Assessment. Agriculture, 16(4), 413. 10.3390/agriculture16040413

European Commission (2020a) Guidance on the approval and low-risk criteria linked to “antimicrobial resistance” applicable to microorganisms used for plant protection in accordance with regulation EC No. 1107/2009. (n.d.).

European Commission (2020b) Guidance on the risk assessment of metabolites produced by microorganisms used as plant protection active substances in accordance with article 77 of Regulation EC No. 1107/2009. (n.d.).

Florensa, A. F., Armenteros, J. J. A., Kaas, R. S., Clausen, P. T. L. C., Nielsen, H., Rost, B., & Aarestrup, F. M. (2026). Whole-genome prediction of bacterial pathogenic capacity on novel bacteria using protein language models with PathogenFinder2. Bioinformatics, btag129. 10.1093/bioinformatics/btag129

Frederiks, C., & Wesseler, J. H. (2019). A comparison of the EU and US regulatory frameworks for the active substance registration of microbial biological control agents. Pest Management Science, 75(1), 87–103. 10.1002/ps.5133

Keller, N. P. (2019). Fungal secondary metabolism: Regulation, function and drug discovery. Nature Reviews. Microbiology, 17(3), 167–180. 10.1038/s41579-018-0121-1

Klassen, J. L., & Currie, C. R. (2012). Gene fragmentation in bacterial draft genomes: Extent, consequences and mitigation. BMC Genomics, 13(1), 14. 10.1186/1471-2164-13-14

Lv, J., Liu, G., Dong, W., Ju, Y., & Sun, Y. (2022). ACDB: An Antibiotic Combination Database. Frontiers in Pharmacology, 13, 869983. 10.3389/fphar.2022.869983

Parks, D. H., Chuvochina, M., Rinke, C., Mussig, A. J., Chaumeil, P.-A., & Hugenholtz, P. (2022). GTDB: an ongoing census of bacterial and archaeal diversity through a phylogenetically consistent, rank-normalized and complete genome-based taxonomy. Nucleic Acids Research, 50, D785–D794.

Poulaki, E. G., & Tjamos, S. E. (2023). Bacillus species: Factories of plant protective volatile organic compounds. Journal of Applied Microbiology, 134(3), xad037. 10.1093/jambio/lxad037

Robin, D. C., & Marchand, P. A. (2019). Evolution of the biocontrol active substances in the framework of the European Pesticide Regulation (EC) No. 1107/2009. Pest Management Science, 75(4), 950–958. 10.1002/ps.5199

Tegenfeldt, F., Kuznetsov, D., Manni, M., Berkeley, M., Zdobnov, E. M., & Kriventseva, E. V. (2025). OrthoDB and BUSCO update: Annotation of orthologs with wider sampling of genomes. Nucleic Acids Research, 53(D1), D516–D522. 10.1093/nar/gkae987

Villaverde, J. J., Sevilla-Morán, B., Sandín-España, P., López-Goti, C., & Alonso-Prados, J. L. (2014). Biopesticides in the framework of the European Pesticide Regulation (EC) No. 1107/2009. Pest Management Science, 70(1), 2–5. 10.1002/ps.3663

Wishart, D. S., Han, S., Saha, S., Oler, E., Peters, H., Grant, J. R., Stothard, P., & Gautam, V. (2023). PHASTEST: faster than PHASTER, better than PHAST. Nucleic Acids Research, 51, W443–W450.

Zhang, S., Merino, N., Okamoto, A., & Gedalanga, P. (2018). Interkingdom microbial consortia mechanisms to guide biotechnological applications. Microbial Biotechnology, 11(5), 833–847. 10.1111/1751-7915.13300

