## Supplementary figures and images for "A new automated pipeline for Whole Genome Shotgun sequencing analysis and hazard characterization of microbial pesticides"

### Supplementary File S8

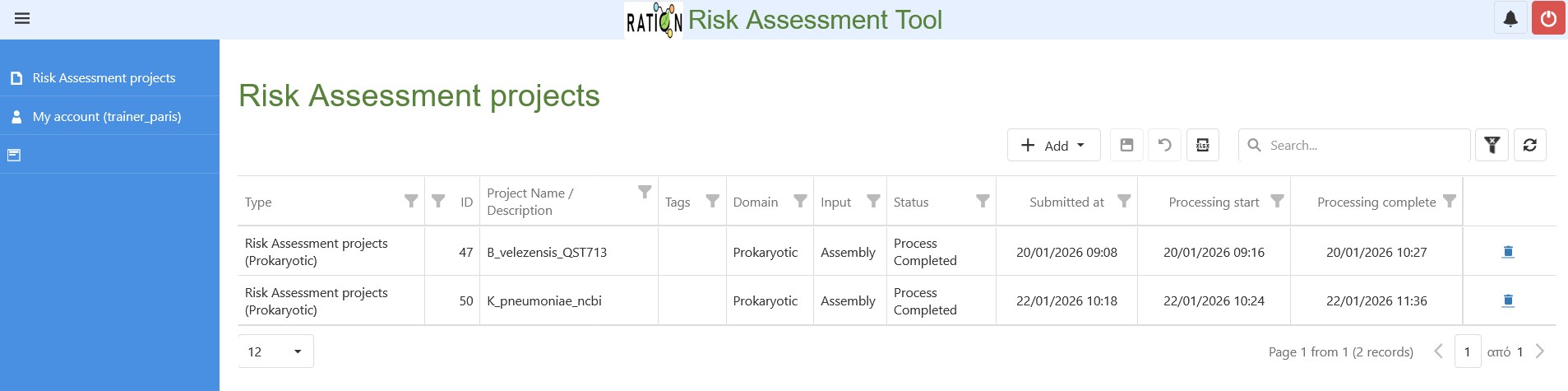

### Supplementary File S9

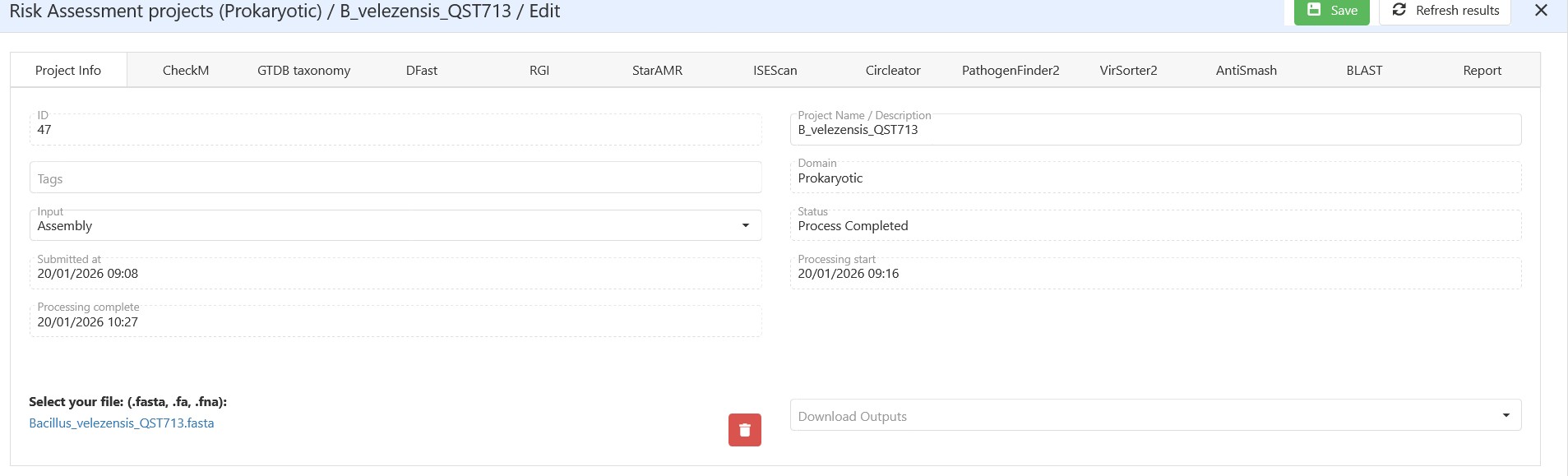
