## Supplementary File S10 for "A new automated pipeline for Whole Genome Shotgun sequencing analysis and hazard characterization of microbial pesticides"

### Tools, Outputs and follow-up procedures:

These guidelines are set up in accordance with the European Food Safety Authority (EFSA) requirements for Whole Genome sequence analysis. (<https://efsa.onlinelibrary.wiley.com/doi/10.2903/j.efsa.2024.8912>).

#### Genome Quality Assessment (CheckM)

CheckM estimates genome completeness and contamination using lineage-specific marker genes. High completeness (>95%) and low contamination (<5%) indicate a reliable genome suitable for downstream risk assessment.

**Follow-up tasks:**

- Verify that completeness is above the expected threshold and contamination is below the accepted threshold for the intended assessment.
- If contamination is elevated, inspect whether the assembly may include mixed strains, contaminant contigs or mis-binned material.
- If completeness is low, treat negative findings, such as absence of AMR genes, mobile elements or biosynthetic clusters, with caution.
- Check whether the genome is closed or highly fragmented, because fragmented assemblies can split AMR loci, mobile elements or biosynthetic gene clusters.
- If the assembly does not meet quality expectations, repeat assembly, sequencing or polishing before drawing risk-relevant conclusions.

For further details on the tool please see:

<https://github.com/Ecogenomics/CheckM.git>

#### Genome Quality Assessment (BUSCO & EukCC)

Both BUSCO and EukCC rely on lineage-specific maker genes to assess completeness and contamination. High completeness (>95%) and low contamination (<5%) indicate a reliable genome suitable for downstream risk assessment.

**Follow-up tasks:**

- Confirm that BUSCO was run with an appropriate lineage dataset for the candidate fungus.
- Compare BUSCO completeness, duplication, fragmentation and missing values; high duplication may indicate haplotigs, heterokaryosis, strain mixture or assembly redundancy.
- Use EukCC contamination estimates to identify possible non-target eukaryotic contamination.
- Treat absence of mycotoxin-related genes or BGCs cautiously when completeness is low or gene prediction quality is uncertain.
- If the fungal genome is highly fragmented, consider whether secondary-metabolite clusters may be split across contigs.

For further details on the tools please see:

<https://github.com/metashot/busco.git>

<https://github.com/EBI-Metagenomics/EukCC.git>

#### Taxonomic Classification (GTDB-Tk) (bacterial)

GTDB-Tk assigns standardized taxonomy based on genome-wide phylogeny and ANI comparisons. High ANI (>95%) to a reference genome supports confident species-level identification.

**Follow-up tasks:**

- Confirm assembly metrics are derived from the final production strain.

Check if species is in a risk class in accordance with e.g., the Technical Rules for Biological Agents (TRBA) of the Federal Institute for Occupational Safety and Health ([BAuA - Technical Rules - Technical Rules for Biological Agents (TRBA) - Federal Institute for Occupational Safety and Health](https://www.baua.de/EN/Service/Technical-rules/TRBA/TRBA)).

For further details on the tool please see:

- <https://github.com/Ecogenomics/GTDBTk.git>

#### Taxonomic Classification (rRNA+ITS) (Fungal)

Barrnap annotates rRNA (5.8S, 18S, and 28S) in your fungal genome. In addition, forty-two is used to predict ITS based on the NCBI ITS database.

**Follow-up tasks:**

- Go to NCBI BLASTn webpage and upload predicted rRNA and ITS sequences.
- Select the “Reference RNA sequences (refseq_rna)” database and “Highly similar sequences (megablast)” option, then run BLAST.
- Compare similarity of each sequence against the database results to determine the most plausible taxonomy

For further details on the tools please see:

<https://github.com/tseemann/barrnap.git>

<https://metacpan.org/pod/FortyTwo::Manual>

#### Annotation (DFAST)

DFAST annotates the assembled genome, identifying CDSs, rRNAs, tRNAs, CRISPRs, GC content, and genome statistics. High N50, single contig assemblies, and high coding ratios indicate a high-quality, near-complete genome.

For further details on the tool please see:

<https://github.com/nigyta/dfast_core.git>

#### Antimicrobial Resistance (RGI, StarAMR)

The EFSA guidelines require the use of 2 independent databases for detection of AMR genes). RGI compares predicted proteins to the **CARD database** to detect AMR genes, reporting identity, coverage, mechanism, and antibiotic class. Hits below regulatory thresholds often represent intrinsic or non-functional homologs. StarAMR (wrapper tool) complements this by predicting resistance phenotypes and MLST profiles using **ResFinder** and PlasmidFinder.

**Follow-up tasks:**

- Assess genomic context of AMR-like hits to confirm absence of mobile genetic elements.
- Confirm phenotypic susceptibility by regulatory guidance.
- Assess whether the AMR determinant is likely intrinsic to the species or acquired, using comparative genomics, taxon-specific literature and public genome searches where possible.

For further details on the tool please see:

<https://github.com/arpcard/rgi.git>

<https://github.com/phac-nml/staramr.git>

#### Mobile Genetic Elements (ISEScan)

ISEScan detects insertion sequences and transposase families. Presence of IS elements suggests genomic plasticity and potential for horizontal gene transfer.

**Follow-up tasks:**

- Map insertion sequences relative to AMR hits, toxin-related genes, virulence-associated genes and BGC boundaries
- Do not interpret the mere presence of IS elements as evidence of AMR transferability; prioritize co-localization and genomic context.

For further details on the tool please see:

<https://github.com/xiezhq/ISEScan.git>

#### Phage & Viral Content (VirSorter2)

VirSorter2 identifies viral and prophage regions. Absence or low-confidence predictions indicate limited viral burden.

**Follow-up tasks:**

- Evaluate the intrinsic/not intrinsic nature of the AMR – follow the guidelines as in the “intrinsic AMR” documentation
- Document lack of co-localization between insertion sequences and risk-associated genes as supporting evidence.
- Retain VirSorter2 and ISEScan outputs as supporting evidence.

For further details on the tool please see:

<https://github.com/jiarong/VirSorter2.git>

#### Pathogenicity (PathogenFinder2)

PathogenFinder2 uses machine learning models to estimate the likelihood a genome originates from a human pathogen (i.e. compares the protein content of the genome against a database of pathogenicity-related protein families). Low probability scores and classification as non-pathogenic support biosafety eligibility.

**Follow-up tasks:**

- Interpret pathogenicity predictions together with taxonomic identity, known biology of the species, literature evidence and the intended product use.
- For positive or borderline predictions, inspect contributing features if available and compare with known virulence-associated functions.

For further details on the tool please see:

<https://github.com/genomicepidemiology/PathogenFinder2.git>

#### Secondary Metabolites & BLAST Analyses (antiSMASH, BLAST)

antiSMASH identifies biosynthetic gene clusters (e.g., lipopeptides like surfactin). BLAST confirms gene identity based on suggested identity/coverage levels and cluster completeness. Presence of intact biocontrol-related clusters supports potential functional efficacy of secondary metabolite production rather than pathogenic risk.

(Results are filtered based on **≥80% sequence identity & ≥70% length coverage)**

**Follow-up tasks (Bacteria):**

- Link detected clusters to peer-reviewed safety (e g. production of toxins like cereulides) and mode-of-action studies (e.g. lipopeptide production).
- Check that metabolites act locally and are not systemically toxic.
- Check potential (amount of) production in product, in situ.
- Evaluate cluster integrity, especially for modular NRPS and PKS pathways that can be split across contigs in draft assemblies.
- Manually inspect neighbouring or overlapping BGCs that may not be resolved as separate regions by antiSMASH.
- Document clusters that require manual curation or additional experimental confirmation in the final interpretation.

**Follow-up tasks (Fungi):**

- Prioritize complete clusters over isolated single-gene hits when interpreting potential mycotoxin production.
- Evaluate whether the candidate species or close relatives are known to produce mycotoxins under relevant growth or formulation conditions.
- For critical mycotoxins, document whether the biosynthetic genes are known and whether the workflow database is expected to cover them.

For further details on how results are generated from the combination of AntiSMASH and BLAST tables please see the FAQs section at <https://agrostis.gr/ration-ra-gui/FAQS>
